# Loss of LasR function leads to decreased repression of *Pseudomonas aeruginosa* PhoB activity at physiological phosphate concentrations

**DOI:** 10.1101/2024.03.27.586856

**Authors:** Amy Conaway, Igor Todorovic, Dallas L. Mould, Deborah A. Hogan

**Author notes:** To whom correspondence should be addressed: Department of Microbiology and Immunology Geisel School of Medicine at Dartmouth Rm 208 Vail Building, Hanover, NH 03755.

## Abstract

While the *Pseudomonas aeruginosa* LasR transcription factor plays a role in quorum sensing (QS) across phylogenetically-distinct lineages, isolates with loss-of-function mutations in *lasR* (LasR– strains) are commonly found in diverse settings including infections where they are associated with worse clinical outcomes. In LasR– strains, the transcription factor RhlR, which is controlled by LasR, can be alternately activated in low inorganic phosphate (Pi) concentrations via the two-component system PhoR-PhoB. Here, we demonstrate a new link between LasR and PhoB in which the absence of LasR increases PhoB activity at physiological Pi concentrations and raises the Pi concentration necessary for PhoB inhibition. PhoB activity was also less repressed by Pi in mutants lacking different QS regulators (RhlR and PqsR) and in mutants lacking genes required for the production of QS-regulated phenazines suggesting that decreased phenazine production was one reason for decreased PhoB repression by Pi in LasR– strains. In addition, the CbrA-CbrB two-component system, which is elevated in LasR– strains, was necessary for reduced PhoB repression by Pi and a Δ*crc* mutant, which lacks the CbrA-CbrB-controlled translational repressor, activated PhoB at higher Pi concentrations than the wild type. The Δ*lasR* mutant had a PhoB-dependent growth advantage in a medium with no added Pi and increased virulence-determinant gene expression in a medium with physiological Pi, in part through reactivation of QS. This work suggests PhoB activity may contribute to the virulence of LasR– *P. aeruginosa* and subsequent clinical outcomes.

**Importance:** Loss-of-function mutations in the gene encoding the *Pseudomonas aeruginosa* quorum sensing (QS) regulator LasR occur frequently and are associated with worse clinical outcomes. We have found that LasR– *P. aeruginosa* have elevated PhoB activity at physiological concentrations of inorganic phosphate (Pi). PhoB activity promotes Pi acquisition as well as the expression of QS and virulence-associated genes. Previous work has shown that PhoB induce RhlR, another QS regulator, in a LasR-mutant in low Pi conditions. Here, we demonstrate a novel relationship wherein LasR represses PhoB activity, in part through the production of phenazines and Crc-mediated translational repression. This work suggests PhoB activity may contribute to the increased virulence of LasR– *P. aeruginosa*.

## Introduction

*Pseudomonas aeruginosa* is a pernicious pathogen that infects burns, nonhealing wounds, eyes, and the airways of people with cystic fibrosis (pwCF) or chronic obstructive pulmonary disease (COPD). *P. aeruginosa* fitness in vivo is due to its ability to establish persistent biofilms and acquire critical nutrients, such as phosphorus, in the host environment. Phosphate is required for cell membranes, nucleic acids, metabolic intermediates, and is used in some signal transduction pathways. The most accessible form of phosphorus is free inorganic phosphate (Pi) which is at ∼1.3 mM in the serum of healthy adults.^(1, 2)^ During infection, however, Pi is restricted by the host as part of nutritional immunity through different mechanisms including the secretion of phosphate-binding proteins.^(3–5)^

Most host phosphorus is in organic forms that require degradation prior to uptake and utilization by microbes.^(6, 7)^ *P. aeruginosa* induces the production of phosphatases, phospholipases, and DNases to access organic phosphate along with high-affinity phosphate transporters in response to low Pi levels via the transcription factor PhoB and its sensor kinase PhoR.^(8–11)^ In *P. aeruginosa*, PhoB also induces the production of specific phenazine small molecules ^(12–16)^ which solubilize phosphate from minerals.^(17)^ PhoR, which activates PhoB through phosphorylation, is regulated by interactions with the high-affinity Pi transporter PstABC via PhoU such that Pi transport inhibits PhoR activation.^(10, 18-25)^ While PhoB is known to positively regulate its own expression ^(26)^, data suggest that PhoB also participates in a negative feedback loop ^(27)^, presumably to limit intracellular Pi concentrations, as has been described in *Escherichia coli*.^(28)^

In addition to the role of the PhoR-PhoB two-component system in Pi acquisition in *P. aeruginosa* and other species such as *E. coli* ^(24, 29)^, *Salmonella enterica* ^(7)^, and *Vibrio cholerae* ^(30)^, these regulators also play diverse roles in the regulation of virulence in multiple pathogens.^(10)^ In *E. coli*, type III secretion system components and effectors are upregulated during Pi limitation in a PhoB-dependent manner.^(31)^ In *V. cholerae*, PhoB inhibits transcription of the genes encoding TcpPH, which positively regulate expression of virulence factors including toxin co-regulated pilus.^(30, 32)^ In *P. aeruginosa,* cross-regulation between PhoR-PhoB and quorum sensing (QS), which contributes to virulence and biofilm formation ^(33–35)^, has been described. *P. aeruginosa* QS is largely controlled by three transcription factors, LasR, RhlR, and PqsR, that are active when QS inducers are at sufficient concentrations, i.e. high cell densities or environments with decreased diffusion. While LasR positively regulates both RhlR and PqsR, many groups have shown that RhlR and PqsR can be activated in the absence of LasR.^(36–39)^ Furthermore, in low Pi conditions, PhoB induces QS ^(14-16, 40, 41)^ and this can occur in a LasR-independent manner.^(13, 42, 43)^ One group of virulence factors induced by both PhoB and QS are phenazines which inhibit immune cell function ^(44, 45)^ and kill other microbes ^(46–50)^ through oxidative stress. Though PhoB binding sites have been identified upstream of some phenazine biosynthesis genes ^(16)^, PhoB induction of phenazine production is largely attributed to increased expression of *rhlR*.^(13, 16, 40)^ Furthermore, *phoR, phoB, rhlR,* and *pqsR* are all required for *P. aeruginosa* phenazine production in co-culture with *Candida albicans* ^(27)^, a fungus that frequently co-infects the airways of pwCF.^(51)^ PhoB regulation of QS suggests low Pi could be a signal for QS remodeling to circumvent reliance on LasR.

*P. aeruginosa* LasR– strains are found in the environment and isolates from both acute and chronic infections like those in the lungs of pwCF where they make up approximately one-third of clinical isolates.^(36, 52-57)^ LasR– isolates are associated with worse disease progression in pwCF ^(57)^ and worse lesions during acute ocular infections.^(54)^ In laboratory evolution experiments, the increased fitness of *P. aeruginosa* LasR– lineages depends on increased CbrA-CbrB activity ^(58, 59)^ which leads to growth advantages in complex media.^(37, 58, 60)^ *P. aeruginosa* CbrA-CbrB is a two-component system that induces expression of the small RNA *crcZ* which sequesters Crc.^(61–64)^ Crc, with Hfq, binds to multiple mRNA targets, many of which encode transporters and catabolic enzymes ^(65)^, and inhibits their translation. CbrA is required for full *P. aeruginosa* virulence ^(66)^.

*P. aeruginosa* virulence regulation is often studied in laboratory media with Pi concentrations that repress PhoR activation of PhoB (e.g. synthetic CF medium (SCFM), 5.1 mM ^(67)^; RPMI medium, 5.6 mM ^(68, 69)^; Luria Broth, 6 mM ^(70)^; or M9 medium, 64 mM ^(70)^). PhoR-PhoB activity is generally studied in media with less than 0.5 mM Pi.^(13, 20, 24, 42)^ These two extremes in Pi concentrations either strongly repress or strongly induce PhoB activity and thus may not be relevant to our understanding of any contributions of PhoB to *P. aeruginosa* virulence at physiological Pi concentrations (1.3 mM Pi in serum ^(1, 4)^). Thus, we aimed to elucidate the relationship between LasR and PhoB at physiologically relevant Pi concentrations. In these studies, we show that LasR– isolates and a Δ*lasR* mutant had elevated PhoR-dependent PhoB activity relative to comparator strains with functional LasR at Pi concentrations in the 0.7 – 1.1 mM range. In contrast, LasR+ and LasR– strains had similar PhoB activity in both low (0.2 mM) and high (10 mM) Pi conditions. Our data demonstrate that a lack of phenazines or reduced Crc activity via CbrA-CbrB led to higher PhoB activity and decreased repression of PhoR-PhoB by Pi. We show that PhoB is required for the Δ*lasR* mutant growth advantages in a medium with no added Pi and the increased expression of multiple virulence factors, including phenazine biosynthetic enzymes and phospholipases, at physiological Pi. This work establishes a novel connection between QS and PhoB wherein LasR represses PhoB activity and PhoB is required for increased virulence gene expression in the Δ*lasR* mutant, in part through reactivation of QS. This model for virulence regulation may aid in understanding why *P. aeruginosa* LasR– strains are associated with poor clinical outcomes.

## Results

### PhoB activity is repressed at lower Pi concentrations in *P. aeruginosa* LasR+ strains than in their *P. aeruginosa* LasR– counterparts

Though PhoB regulates virulence factor production across species ^(10, 30, 31)^, there are only a limited number of in vitro studies on PhoB activity at Pi concentrations similar to those found in human hosts. Thus, we sought to determine the Pi concentrations necessary to repress PhoB activity. In *P. aeruginosa* strain PA14, PhoB activity was monitored by assessing the activity of alkaline phosphatase (AP), which is encoded by the PhoB-regulated gene *phoA* ^(8, 71, 72)^ in colonies on agar containing the colorimetric AP substrate bromo-4-chloro-3-indolyl phosphate (BCIP) ^(9, 73)^ (Fig. 1A). Using gradient plates with range of Pi concentrations (0.1 mM - 1 mM) ^(27)^ in MOPS-glucose medium ^(74)^, we found that 0.7 mM Pi repressed AP activity in the wild type. Interestingly, a Δ*lasR* mutant had AP activity across the entire gradient with a reduction in activity at higher Pi concentrations (Fig. 1A). Consistent with PhoB regulation of AP activity, the Δ*phoB* mutant had no detectable AP activity at any concentration, and the Δ*pstB* mutant, which has constitutive PhoB activity due to de-repression of PhoR ^(27, 75)^, had strong AP production at all Pi concentrations tested (Fig. 1A). At 0.7 mM Pi, these was AP activity in colonies of the Δ*lasR* mutant, but not the wild type colonies or those formed by the Δ*lasR* mutant complemented with *lasR* at the native locus (Fig. 1B). As expected, neither the Δ*lasRΔphoB* or Δ*lasRΔphoR* mutants had AP activity (Fig. 1B). Furthermore, a clinical isolate from a corneal infection, DH2590 (262K in Hammond, *et al.* ^(54)^), which has a loss-of-function (LOF) mutation in *lasR,* encoding the non-functional LasR^I215S^ variant, also expressed AP at 0.7 mM Pi and AP activity was reduced by replacing the endogenous *lasR* allele with a functional PA14 *lasR* allele at the native locus (Fig. 1B). In addition, we compared AP activity in two clinical isolate pairs (Fig. 1C) from CF sputum samples wherein one isolate has a LOF mutation in *lasR* and the other does not.^(76)^ In both cases, the LasR– isolates had more AP activity than their LasR+ counterparts at 0.7 mM Pi.

**Fig. 1.**
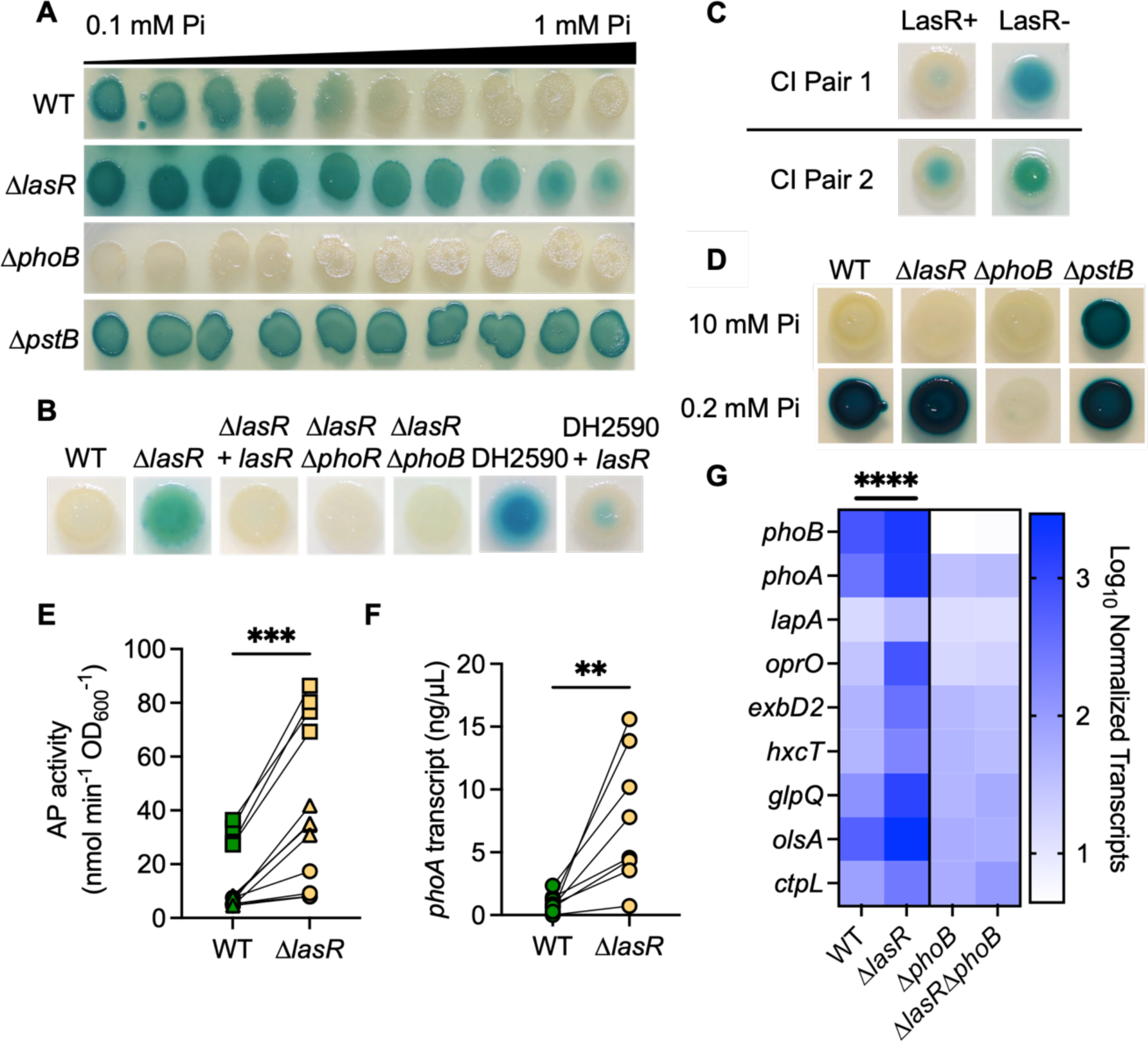
*P. aeruginosa* LasR– strains require higher Pi to repress PhoB activity. **A)** *P. aeruginosa* wild type (WT), Δ*lasR*, Δ*phoB* and Δ*pstB* were spotted on gradient plates of MOPS agar with a range of Pi concentrations (0.1 - 1 mM) and 60 µg/mL BCIP to indicate alkaline phosphatase (AP) activity. **B)** *P. aeruginosa* strains, including DH2590 (a LasR– clinical isolate (CI) and a derivative in which the *lasR* allele was replaced with PA14 *lasR*) on MOPS agar with BCIP and 0.7 mM Pi. **C)** *P. aeruginosa* CI LasR+/LasR– pairs from the sputum of two pwCF on MOPS agar with BCIP and 0.7 mM Pi. **D)** Colony biofilms on MOPS agar with 0.2% glucose and either 0.2 or 10 mM Pi and BCIP. For **A-D**, similar results were obtained in three replicate experiments; a representative experiment is shown. For panels **E-G**, *P. aeruginosa* was grown as colony biofilms on MOPS agar with 0.7 mM Pi. **E)** AP activity in WT and Δ*lasR* after 12 h at 37 °C. Data from replicates collected on the same day have the same shape. Data analyzed using a paired, two-tailed t-test (n = 12). **F)** *phoA* transcripts in WT and Δ*lasR* were measured by qRT-PCR on different days and normalized to the housekeeping gene transcript *ppiD*. Data analyzed using a paired, two-tailed t-test (n = 8). **G)** Levels of PhoB-controlled transcripts and *phoB* itself were assessed using NanoString multiplex technology. Nine PhoB-regulated transcripts are shown as normalized counts. Data were analyzed using a two-way ANOVA; significant differences between WT-Δ*lasR,* WT-Δ*phoB*, and *ΔlasR-*Δ*lasRΔphoB* (p < 0.0001, n = 2-3). There were no significant differences between the Δ*phoB-*Δ*lasRΔphoB* mutants (p = 0.46, n = 3). Statistical differences for each transcript are available in Supplemental File 1. For all panels, asterisks denote significance (p ≤ 0.01 = **\*\***, p ≤ 0.001 = **\*\*\***, p ≤ 0.0001 = **\*\*\*\***).

Both the Δ*lasR* mutant and the wild type showed PhoB activity at lower Pi concentrations (< 0.7 mM). We found that at 0.2 mM Pi, the concentration frequently used to study PhoB activity ^(13, 14, 20, 24, 77)^, the wild type and Δ*lasR* strains had similar AP activity (Fig. 1D). Additionally, neither strain showed AP production at 10 mM Pi (Fig. 1D) and no AP activity was observed in either the wild-type or the Δ*lasR* strain on LB agar, which is reported to have 6 mM Pi ^(70)^ (Fig. S1A). The Δ*phoB* mutant did not have AP activity even at 0.2 mM Pi and the Δ*pstB* mutant had AP production at 0.2 mM Pi, 10 mM Pi, and on LB. To quantify differences AP activity, we used the soluble colorimetric substrate p-nitrophenyl phosphate (PNPP). The Δ*lasR* mutant had significantly more AP activity than the wild type at 0.7 mM Pi (Fig. 1E) while growth was similar (Fig. S1B). Similarly, qRT-PCR analysis of *phoA*, which encodes AP, found significantly higher transcript levels in the Δ*lasR* mutant compared to the wild type (Fig. 1F).

To directly assess the levels of transcripts of other genes within the PhoB regulon, we utilized NanoString multiplex technology as previously published ^(27)^. At 0.7 mM Pi, the Δ*lasR* mutant had significantly higher levels of PhoB-regulated transcripts including *phoB* itself and genes encoding phosphatases (*phoA* and *lapA*), a phosphate transporter (*oprO*), a type two secretion system protein required for secretion of LapA (*hxcT*) ^(78)^, a putative TonB transporter (*exbD2*) ^(79)^, a phosphodiesterase (*glpQ*), a low-phosphate ornithine lipid biosynthetic protein (*olsA*) ^(80)^, and a Pi chemotaxis protein (*ctpL*).^(81)^ None of these transcripts were significantly different when the Δ*phoB* and Δ*lasRΔphoB* mutants were compared (Fig. 1G and Supplemental File 1).

### Other QS mutants and a mutant lacking QS-controlled phenazines have active PhoB at higher Pi concentrations than the wild type

LasR positively regulates other QS transcription factors including RhlR and PqsR and some Δ*lasR* mutant phenotypes are due to their decreased activity. However, RhlR and PqsR are capable of inducing QS-controlled genes in the absence of LasR in specific strains and conditions.^(36, 37, 59, 82)^ Thus, we assessed the role of RhlR and PqsR in the repression of PhoB. As shown in Fig. 2A, AP activity was significantly elevated in the *ΔrhlR* and *ΔpqsR* mutants, as in the Δ*lasR* mutant, relative to the wild type at 0.7 mM Pi. Across a gradient plate of 0.5 – 1.5 mM Pi, AP activity in all QS mutants was inhibited at a significantly higher Pi concentration than the wild type (Fig. 2B). While AP activity in the Δ*rhlR* and Δ*pqsR* strains was inhibited by Pi > 0.9 mM, activity in the LasR— strains (Δ*lasR,* Δ*lasRΔrhlR,* and Δ*lasRΔpqsR*) was inhibited by Pi > 1 mM (Fig. 2B).

**Fig. 2.**
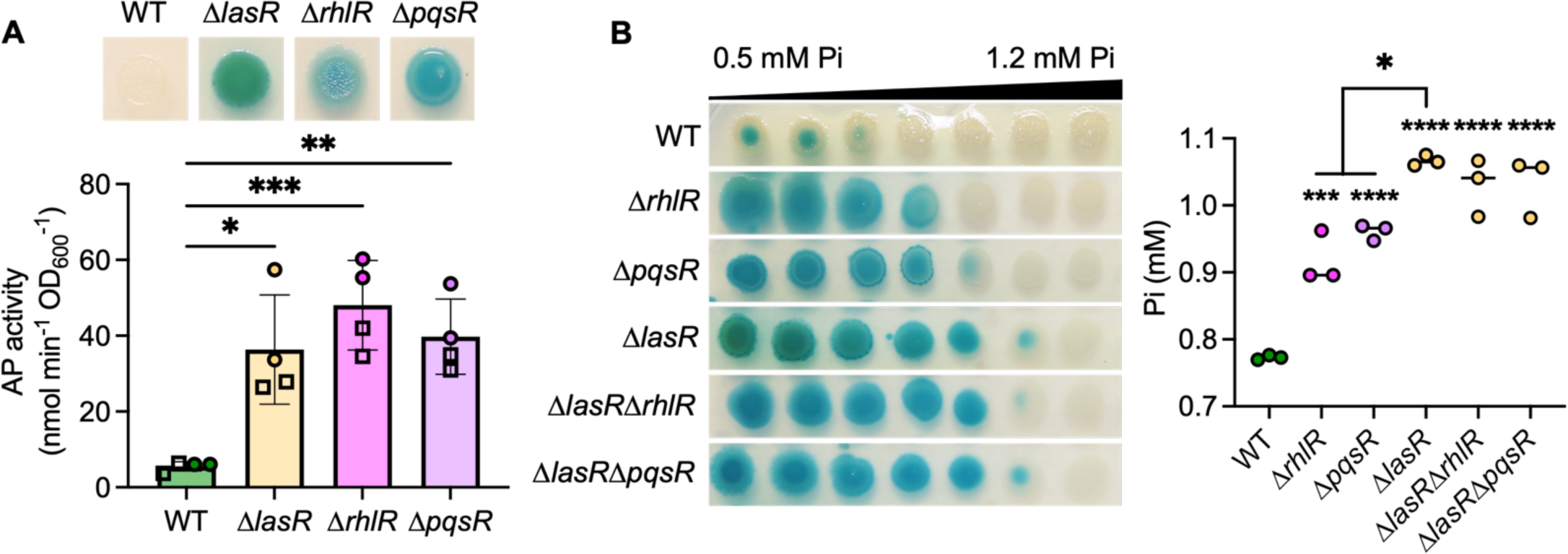
*P. aeruginosa* quorum sensing (QS) mutants have active PhoB at higher Pi than the wild type. **A)** Colony biofilms of wild type (WT) and the indicated QS mutants (Δ*lasR*, Δ*rhlR*, Δ*pqsR*) were grown on MOPS agar with 0.7 mM Pi and BCIP (top) or on medium without BCIP for analysis of AP activity using the colorimetric PNPP substrate. Data from replicates collected on the same day have the same shape. Data analyzed by an ordinary one-way ANOVA with Tukey’s multiple comparisons tests (n = 4). **B)** *P. aeruginosa* was grown on a plate with a gradient of Pi concentrations, with 0.5 to 1.2 mM shown. Quantitative data analyzed using a one-way ANOVA (n = 3) (asterisks denote significance from the wild type). For all panels, asterisks denote significance (p ≤ 0.05 = **\***, p ≤ 0.01 = **\*\***, p ≤ 0.001 = **\*\*\***, p ≤ 0.0001 = **\*\*\*\***).

As each of the QS transcription factor mutants, Δ*lasR*, Δ*rhlR*, and Δ*pqsR*, produce fewer phenazines than the wild type in late exponential and early stationary phase cultures ^(83, 84)^, we tested whether the absence of phenazines was sufficient to increase PhoB activity at moderate Pi concentrations. *P. aeruginosa* produces multiple phenazines including phenazine-1-carboxylic acid (PCA and the PCA derivatives 5-methyl-PCA (5MPCA), pyocyanin (PYO), and phenazine-1-carboxamide. PCA is synthesized by proteins encoded by two highly similar operons, *phzA1-G1* and *phzA2-G2.* The mutant lacking both *phz* operons (Δ*phzA1-G1ΔphzA2-G2*), referred to as Δ*phz*, had significantly elevated AP activity compared to the wild type at 0.7 mM Pi (Fig. 3A). The Δ*phz1* mutant (Δ*phzA1-G1*) was not different from the wild type and deletion of either of the adjacent phenazine-modifying genes required for 5MPCA and PYO biosynthesis, *phzM* or *phzS*, did not lead to changes in AP production at 0.7 mM (Fig. S2A). In contrast, the Δ*phz2* mutant (Δ*phzA2-G2*) phenocopied Δ*phz* and had significantly more AP production than the Δ*phz1* mutant or the wild type (Fig. 3A). We and others have previously published data identifying *phzA2-G2* as the predominant contributor of PCA.^(27, 84-86)^

**Fig. 3.**
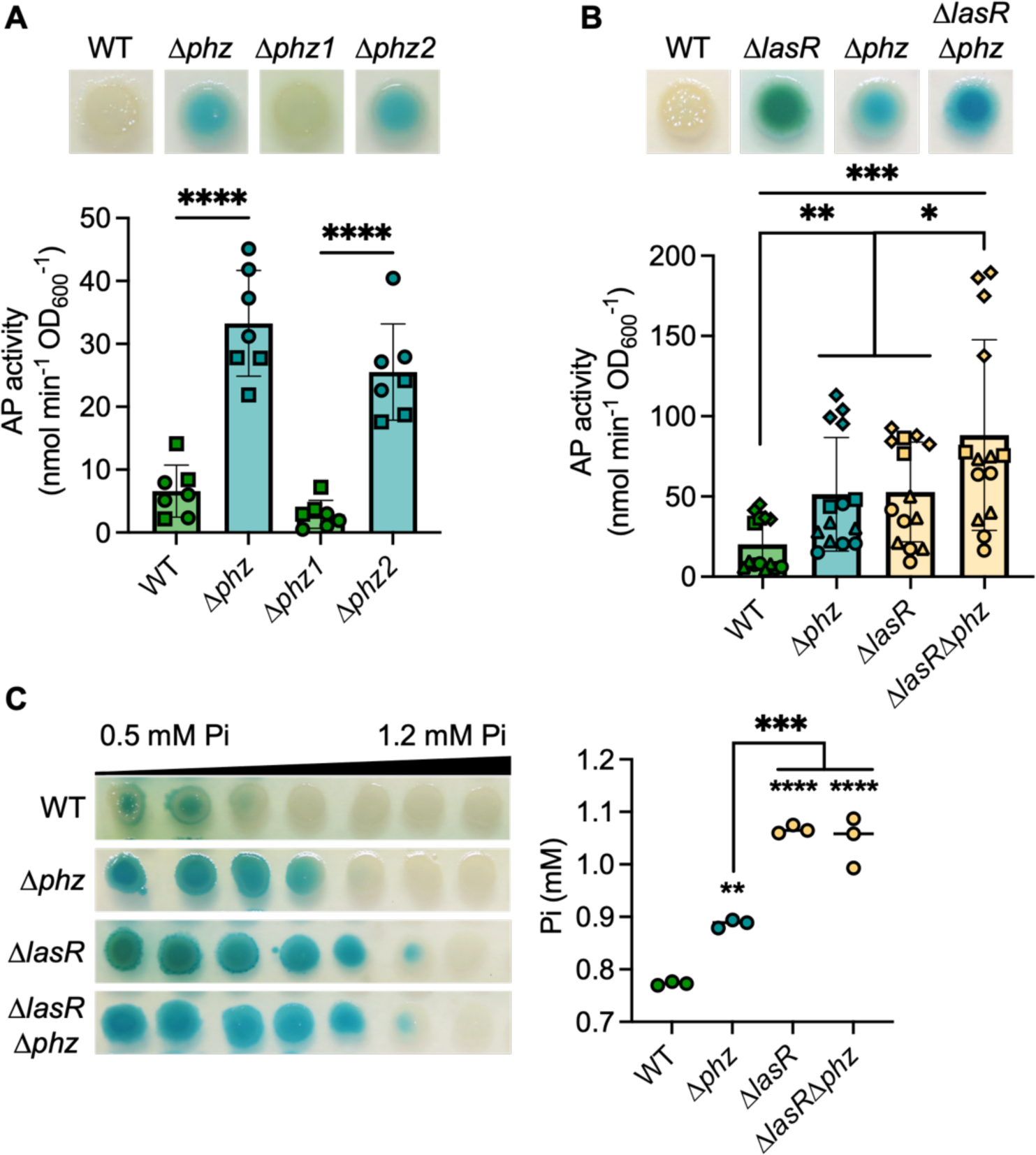
The loss of phenazines promotes PhoB activity. **A)** AP activity in wild type (WT) and mutants Δ*phz1*, Δ*phz2*, and Δ*phz* (lacking *phz1* and *phz2* operons) grown on MOPS agar with 0.7 mM Pi and BCIP (top) or on medium without BCIP for analysis of AP activity using the colorimetric PNPP substrate. Data from replicates collected on the same day have the same shape. Data were analyzed using a one-way ANOVA and Tukey’s multiple comparisons test (n = 7). **B)** Analysis of WT and the Δ*lasR*, Δ*phz*, and Δ*lasR*Δ*phz* mutants. The *ΔlasRΔphz* mutant had significantly more AP activity than the *ΔlasR* and Δ*phz* strains. There are no significant differences between Δ*lasR* and Δ*phz* (p=0.99) (n = 12). **C)** *P. aeruginosa* was grown on plates with a gradient of Pi; the average concentration of Pi that inhibits AP activity is graphed to the right (n = 3). Data analyzed using a one-way ANOVA; asterisks denote significance from the wild type. Asterisks denote significance (p ≤ 0.05 = **\***, p ≤ 0.01 = **\*\***, p ≤ 0.001 = **\*\*\***, p ≤ 0.0001 = **\*\*\*\***).

We found no difference in AP activity at 0.7 mM Pi between the Δ*lasR* and Δ*phz* mutants (Fig. 3B). However, a *ΔlasRΔphz* mutant had significantly elevated AP activity compared to both single mutants (Fig. 3B). Both the Δ*lasR* and *ΔlasRΔphz* mutants had detectable AP activity at significantly higher Pi concentrations than Δ*phz* (Fig. 3C) suggesting that PhoB activity was derepressed by an additional phenazine-independent mechanism in the Δ*lasR* mutant. Consistent with this model, the Δ*phz* mutant was repressed by a similar Pi concentration as the Δ*rhlR* and Δ*pqsR* mutants, but not their LasR– counterparts (Fig. 2B).

### CbrA-CbrB-Crc impacts PhoB activity in the Δ*lasR* mutant

Several studies suggest that *P. aeruginosa* strains lacking LasR activity have elevated CbrA-CbrB activity in many conditions.^(58, 60)^ CbrA-CbrB activity leads to relief of Crc-mediated translational repression of diverse mRNA targets including many that encode catabolic enzymes and transporters ^(64, 65, 87)^ (Fig. 4A). In both BCIP and PNPP assays, we found that in a Δ*lasRΔcbrB* mutant did not have AP activity at 0.7 mM Pi and that AP production was restored by complementation of *cbrB* at the native locus (Fig 4B). PhoB activity in both the Δ*cbrB* and Δ*lasRΔcbrB* mutants was similarly repressed when Pi > 3 mM (Fig. S2B). Additionally, AP activity in colonies grown on medium with 0.7 mM Pi was restored to a Δ*lasRΔcbrB* mutant by deletion of *crc* (Fig. 4B) suggesting that the increase in Pi repression of PhoB in the Δ*lasRΔcbrB* is due to elevated Crc activity (Fig. 4A for pathway). A Δ*crc* single mutant also had significantly elevated AP activity compared to the wild type and *crc*-complemented strain (Fig. 4C). There was no difference in AP activity between the Δ*lasR* and Δ*lasRΔcrc* mutants at 0.7 mM Pi, though both were significantly elevated compared to the Δ*crc* mutant (Fig. 4D). Additionally, the Δ*lasR* and Δ*lasRΔcrc* mutants both showed significantly decreased PhoB sensitivity to Pi than the Δ*crc* mutant (Fig. 4E).

**Fig. 4.**
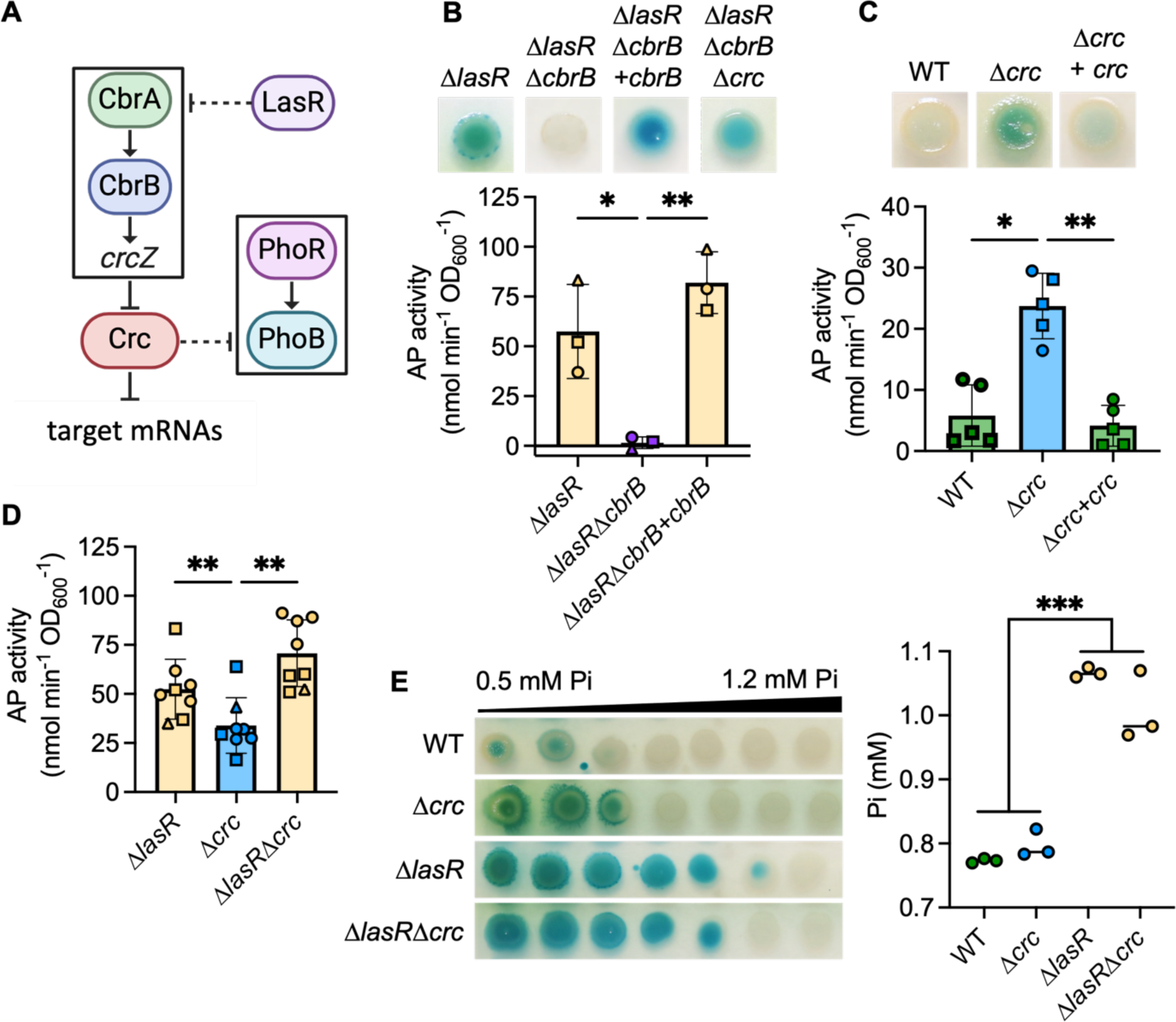
The CbrA-CbrB-Crc pathway promotes PhoR-PhoB activity in LasR– strains. **A)** A proposed model of the relationship between the CbrA-CbrB and PhoR-PhoB two component systems; *crcZ*, a small RNA; Crc, which acts as a translational repressor in complex with Hfq. For figures **B** and **C**, AP activity in indicated strains grown on MOPS agar with 0.7 mM Pi and BCIP (top) or on medium without BCIP for analysis of AP activity using the colorimetric PNPP substrate. Data from replicates collected on the same day have the same shape. Data were analyzed using a one-way ANOVA and Tukey’s multiple comparisons test. **B)** AP activity in Δ*lasR*, Δ*lasRΔ*cbrB, Δ*lasRΔcbrB*+cbrB, and Δ*lasRΔcbrB+crc* mutants (n = 3). **C)** AP activity in the wild type (WT), Δ*crc* mutant and its complemented derivative (n = 5). **D)** AP activity in Δ*lasR*, Δ*crc*, and Δ*lasR*Δ*crc* (n = 8). **E)** *P. aeruginosa* was grown on plates with a gradient of Pi (0.5 – 1.5 mM Pi) and BCIP. The average concentration of Pi which inhibited AP activity is graphed to the right (n = 3). Asterisks denote significance (p ≤ 0.05 = **\***, p ≤ 0.01 = **\*\***, p ≤ 0.001 = **\*\*\***).

### PhoB activity in the *P. aeruginosa* Δ*lasR* strain provides fitness advantages when Pi is limited

To determine if the increased PhoB activity in Δ*lasR* cells increased fitness upon depletion of Pi, we grew the wild type, the Δ*lasR* mutant, and their respective Δ*phoB* derivatives overnight in LB followed by sub-culture into MOPS-glucose medium with no added Pi (Fig. 5A). The wild type and the Δ*lasR* mutant showed similar initial growth rates. However, in the post-exponential phase, the Δ*lasR* mutant reached significantly higher densities than the wild type. Both the Δ*phoB* and Δ*lasRΔphoB* mutants showed minimal growth in the no added Pi media and were not significantly different from each other.

**Fig. 5.**
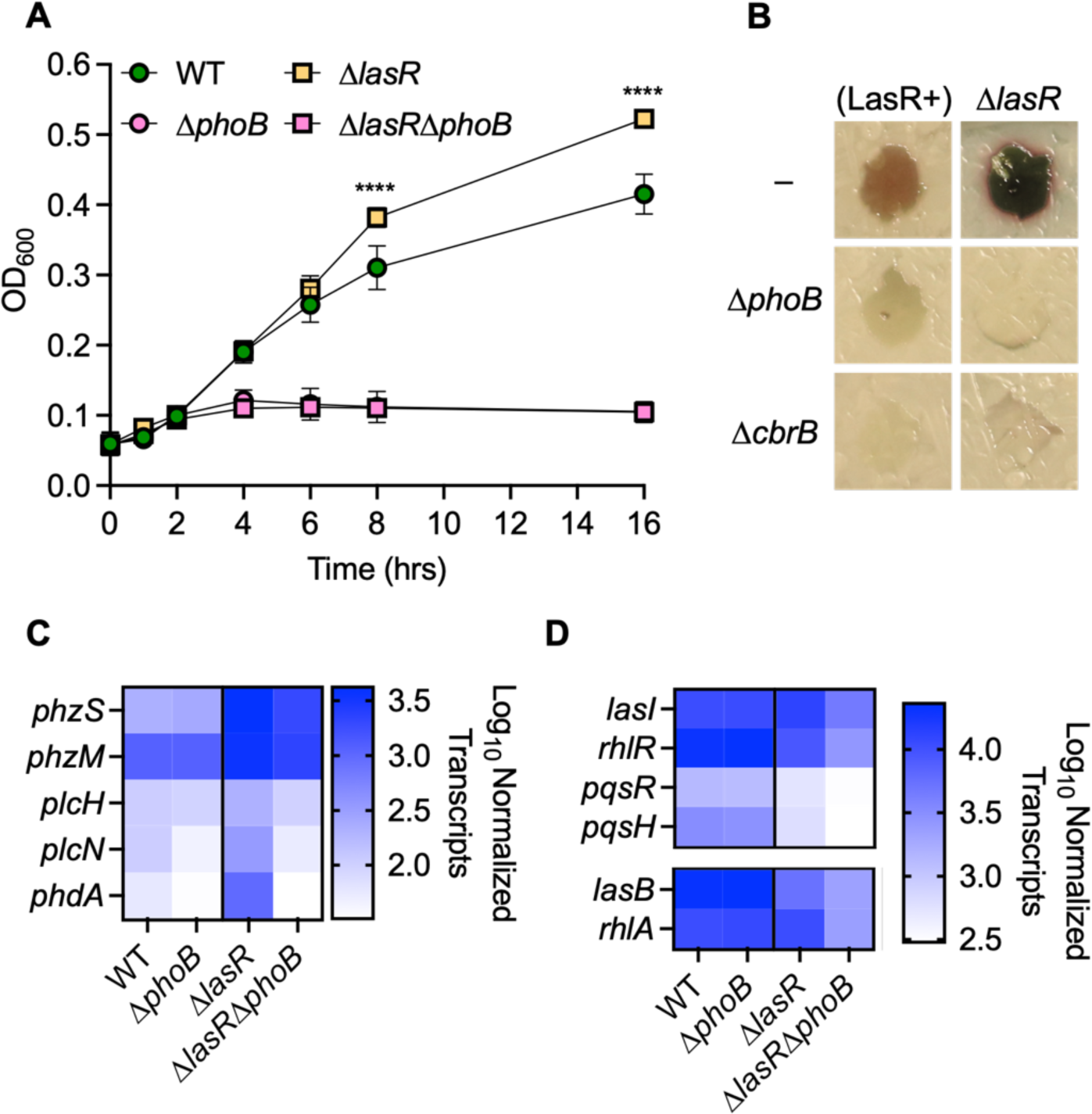
PhoB activity mediates fitness advantages and antagonism in Δ*lasR P. aeruginosa*. **A)** *P. aeruginosa* wild type (WT), Δ*lasR*, Δ*phoB*, and Δ*lasR*Δ*phoB* were grown in MOPS liquid medium with no added Pi at 37 °C. Data analyzed using a two-way ANOVA with Tukey’s multiple comparisons test (n = 3). Asterisks denote significance (p ≤ 0.0001 = **\*\*\*\***). **B)** *P. aeruginosa* colonies were grown on a lawn of *Candida albicans*. Wild-type (LasR+) *P. aeruginosa* produces 5MPCA which is converted to red pigment inside *C. albicans*. The Δ*lasR* mutant produced more 5MPCA than the wild type. 5MPCA production required PhoB and CbrB in both strains (n = 3). **C)** NanoString analysis of PhoB-regulated transcripts shown as Log_10_ normalized counts (n = 2 - 3). All transcripts shown were significantly higher in the *ΔlasR* mutant compared to both the wild type (p ≤ 0.0001) and the Δ*lasRΔphoB* mutant (p ≤ 0.0016). **D)** PhoB contribution to regulation of QS genes by NanoString analysis of QS-regulated transcripts between the WT or Δ*phoB* mutant (p > 0.30) and the Δ*lasR* and Δ*lasR*Δ*phoB*. All transcripts were significantly lower in the Δ*lasRΔphoB* mutant than the Δ*lasR* mutant (p ≤ 0.01)*. lasB, rhlR, pqsR,* and *pqsH* were significantly lower in the Δ*lasR* mutant than the wild type (p < 0.0001). Complete NanoString dataset available in Supplemental File 1.

### PhoB activity mediates the expression of virulence determinants in LasR– *P. aeruginosa,* in part through QS

Multiple studies have shown that there are alternative mechanisms for the activation of RhlR in LasR– strains ^(13, 36-38, 42, 43)^ and that phosphate limitation promotes RhlR activity.^(13-16, 42, 43)^ We have previously published that both PhoB and RhlR are required for 5MPCA production by *P. aeruginosa* resulting in antagonism of *C. albicans* in co-culture.^(27)^ A derivative of 5MPCA accumulates inside *C. albicans* cells as a red pigment. Here we show evidence of increased 5MCPA production by the Δ*lasR* mutant in co-culture with *C. albicans.* 5MPCA production by both LasR+ and LasR– *P. aeruginosa* required PhoB and CbrB (Fig. 5B).

The NanoString codeset used in Fig. 1G included genes that encode proteins associated with phosphate acquisition, virulence, and QS.^(27)^ We analyzed the abundance of multiple transcripts encoding virulence-associated proteins, including phenazine biosynthetic enzymes (*phzM* and *phzS*), phospholipases (*plcH* and *plcN*), and an exopolysaccharide matrix regulator (*phdA*) ^(88)^ in the wild type and Δ*lasR, ΔphoB,* and Δ*lasRΔphoB* mutants. These virulence-associated transcripts were significantly higher in the Δ*lasR* mutant compared to the wild type (Fig. 5C and Supplemental File 1). Three of these transcripts have also been identified as part of the PhoB regulon (*plcH* ^(89)^, *plcN* ^(12, 90, 91)^ and *phdA* ^(79, 92)^) while others have not (*phzM* and *phzS*). However, all five transcripts were significantly lower in the Δ*lasRΔphoB* mutant compared to the Δ*lasR* strain and no differences were detected when the Δ*phoB* and Δ*lasR*Δ*phoB* mutants were compared. While *plcN* and *phdA* were significantly lower in Δ*phoB* compared to the wild type, *phzM, phzS,* and *plcH*, were only PhoB-dependent in the Δ*lasR* mutant. We then sought to determine if PhoB contributes to QS. We found significantly higher abundance of transcripts encoding proteins involved in QS regulation (*rhlR* and *pqsR*) and autoinducer synthesis (*lasI* and *pqsH*) as well as transcripts that can be induced by RhlR (*rhlA* and *lasB*) in the Δ*lasR* mutant compared to the Δ*lasRΔphoB* mutant (Fig. 5D and Supplemental File 1). As expected, the *lasB, rhlR, pqsR*, and *pqsH* transcripts were also significantly lower in the Δ*lasR* mutant compared to the wild type. Consistent with the model that PhoB is not active in the wild type at 0.7 mM Pi, there were no significant differences in QS-related transcript abundance between the Δ*phoB* mutant and the wild type.

## Discussion

In this manuscript, we showed that LasR– *P. aeruginosa* laboratory strains and clinical isolates had elevated PhoB-dependent AP activity and increased expression of the PhoB regulon at 0.7 mM Pi. We found that the range of PhoB-permissive Pi concentrations was broader for LasR– *P. aeruginosa*, though PhoB was still repressed at high Pi concentrations. Importantly, these intermediate Pi concentrations where LasR–*P. aeruginosa* showed more PhoB activity are similar to those found in the serum of healthy adults.^(1, 2)^ We also found that elevated PhoB activity was common across QS mutants and this was possibly due to reduced phenazine production. However, LasR– *P. aeruginosa* had active PhoB at even higher Pi concentrations than the other QS mutants, suggesting differences in PhoB regulation.

LasR– *P. aeruginosa* are more fit than their LasR+ counterparts in many contexts. LasR– *P. aeruginosa* are known to have elevated CbrA-CbrB activity, conferring growth advantages in complex media through decreased Crc activity.^(58, 60)^ Other studies have shown that LasR– *P. aeruginosa* are more resistant to cell lysis at high pH ^(93)^ and more fit in microoxic conditions and normoxically grown colony biofilms.^(94)^ Future studies are necessary to determine if these phenotypes are interrelated. We showed that CbrB was required for increased PhoB activation in LasR– *P. aeruginosa* and a *crc* mutation was sufficient to induce PhoB activity, even in a LasR+ strain. Importantly, the Δ*crc* mutant had less AP activity and was inhibited by lower Pi concentrations than the Δ*lasR* mutant. These data suggest Crc– *P. aeruginosa* do not phenocopy LasR– strains.

The elevated PhoB activity in the Δ*lasR* mutant contributed to fitness upon a shift from LB to a medium without added Pi. As phosphate accessibility can be transient inside the host, increased PhoB activity may be beneficial in building up stores of polyphosphate for utilization when cells experience low Pi stress. In addition to regulating phosphate acquisition, PhoB activity has also been shown to promote RhlR expression and subsequent PYO production, swarming motility, and cytotoxicity.^(13, 15, 42, 43, 90)^ Here we showed that the Δ*lasR* mutant had PhoB-dependent increased expression of virulence-associated genes including phenazine biosynthetic enzymes and phospholipases like PlcH. PlcH is known to degrade airway surfactant and we have previously shown that its expression leads to a decline in lung function.^(95)^ We also showed PhoB and CbrB are both required for increased production of the antifungal phenazine 5MPCA by the Δ*lasR* mutant in co-culture with *C. albicans*.

The roles for phenazines in PhoB regulation are complex. As all QS mutants produce fewer phenazines in late exponential phase and early stationary phase ^(83, 84)^, we proposed phenazines may repress PhoB activity. We demonstrated that phenazine-deficient *P. aeruginosa* had similarly elevated PhoB activity to the single QS mutants. Unlike the Δ*rhlR* and Δ*pqsR* mutants, LasR– *P. aeruginosa* can still produce phenazines in certain contexts.^(13, 36-38)^ We observed that the Δ*lasR* mutant appeared greener on BCIP agar at 0.7 mM Pi which we speculate is due to the production of the blue-green phenazine PYO as this was not observed in the phenazine-deficient Δ*lasRΔphz* mutant (Fig. 3B). Thus, the higher AP activity in the Δ*lasR*Δ*phz* compared to the Δ*lasR* may be due to the absence of phenazines that reduce PhoB activation at moderate Pi concentrations. It is also of note that PhoB activity promotes phenazine production, potentially through direct binding upstream of the *phz* operons ^(16)^, but more likely through RhlR activity.^(13, 42, 43)^ This increased phenazine production could then inhibit further PhoB activity, providing negative feedback. However, our data suggests PCA, not PYO or 5MPCA, inhibits PhoB as loss of the biosynthetic enzymes PhzM or PhzS did not induce PhoB activity. It cannot be assumed that cells producing visible PYO also have high PCA and reduced PhoB activity. An example of this is the Δ*crc* mutant which we showed had elevated AP activity and is known to produce more PYO through de-repression of PhzM.^(96)^ In some instances, increased PCA conversion to PYO could result in PCA depletion and thus aid in the activation of PhoB. Further work is needed to fully understand the interplay between PhoB activity and phenazines.

It is well established that in permissive Pi conditions, PhoB can promote the expression QS genes directly ^(40)^ or through RhlR ^(15, 16, 43, 90)^, allowing cells to circumvent reliance on LasR as a QS regulator.^(13, 42)^ Our data support this existing model by showing QS gene expression is PhoB-dependent in a Δ*lasR* mutant but not the wild type at 0.7 mM Pi. Importantly, we have shown increased PhoB activity in LasR– *P. aeruginosa* compared to LasR+ strains and this has not been identified in previous publications. The relatively low (0.4 – 0.5 mM) and high (4 – 4.5 mM) Pi concentrations used in the past likely hindered the observation of increased PhoB activity in LasR– *P. aeruginosa* and demonstrates the utility of gradient agar plates. As we found PhoB is active in the Δ*lasR* mutant at higher Pi concentrations than the wild type, the previously reported PhoB-driven reorganization of QS may occur in LasR– *P. aeruginosa* under conditions that otherwise repress PhoB activity in LasR+ strains.

Further work is needed to elucidate the mechanisms by which LasR activity represses PhoB activity at intermediate Pi concentrations. There is evidence that PhoB can be spontaneously phosphorylated by acetyl phosphate in vitro ^(97)^ but the concentrations required for this reaction suggest it is unlikely to occur inside cells.^(98)^ Therefore, we expect changes in PhoB activity are mediated by its sensor kinase PhoR. PhoR activity is suppressed by interactions with the Pi transporter PstABC and PhoU.^(18, 21, 23)^ Thus, changes in expression of these proteins could alter PhoR-PhoB activity. We observed no differences in transcript abundance of *phoU* or *pstA* between the wild type and Δ*lasR* mutant (Supplemental File 1) and so believe this model is unlikely. However, other changes in Pi transport into the cell could still contribute to differences in PhoB activity. While the mechanisms of PhoR-PhoU-PstABC interactions are not well understood, PhoR has been shown to interact with PhoU through its Per-Arnt-Sim (PAS) domain.^(23)^ PAS domains in other organisms bind a broad range of ligands ^(99)^ and can modify kinase activity.^(100, 101)^ Thus, we propose that changes to the intracellular environment in phenazine-deficient cells or cells with low Crc activity may stimulate PhoR activity through its PAS domain. Future work into the mechanisms of the PhoR PAS domain would also be beneficial for understanding similar kinases in *P. aeruginosa* and other organisms as PAS domains are abundant but remain poorly understood.^(102)^ Understanding LasR– *P. aeruginosa* virulence in physiologically relevant conditions is critical as LasR– isolates are frequently found in both acute and chronic infections and these findings may be relevant to *P. aeruginosa* when in settings that do not induce QS regulation. We propose that increased PhoB-mediated QS and virulence gene expression in LasR– *P. aeruginosa* may contribute to their association with worse outcomes.^(54, 57)^

In conclusion, our data support a novel model where LasR represses PhoB activity and thus LasR– *P. aeruginosa* have elevated PhoB activity at physiological Pi concentrations. Others have shown that PhoB activity can induce RhlR activity in the absence of LasR, leading to increased production of virulence factors like PYO.^(13, 42)^ Here we have shown QS and virulence-related gene expression in the Δ*lasR* mutant is highly PhoB dependent at a physiologically relevant Pi concentrations.

## Materials and Methods

### Strains and growth conditions

Bacterial strains and plasmids used in this study are listed in **Table S1**. Bacteria were streaked from frozen stocks onto LB (lysogeny broth) with 1.5% agar.^(103)^ Planktonic cultures were grown in 5 mL LB medium in 18 mm Borosilicate glass tubes on a roller drum at 37 °C in the indicated medium.

### Construction of in-frame deletion, complementation, and expression plasmids

Construction of in-frame deletion and complementation constructs was performed using yeast cloning techniques in *Saccharomyces cerevisiae* as previously described ^(103)^ or Gibson assembly.^(104, 105)^ In-frame deletion and chromosomal complementation constructs were made using the allelic replacement vector pMQ30.^(106)^ Plasmids were purified from yeast using Zymoprep Yeast Plasmid Miniprep II according to the manufacturer’s protocol and transformed into *E*. *coli* strain S17/λpir by electroporation. Plasmids were introduced into *P*. *aeruginosa* by conjugation and recombinants were obtained using sucrose counter-selection. Genotypes were screened by PCR and plasmid constructs were confirmed by sequencing.

### Alkaline phosphatase activity assessment

Agar plates were made as previously described ^(27)^ with MOPS minimal medium ^(74)^ with 0.7 mM K_2_HPO_4_/KH_2_PO_4_, 60 µg/mL BCIP (Sigma Aldrich # 11383221001), 20 mM glucose and 15 g/L agar (referred to here as MOPS agar). Overnight cultures of *P. aeruginosa* were grown in LB, normalized to an OD_600_ of 1, then 5 µL were spotted on MOPS agar and incubated at 37 °C for 24 h. Plates with a gradient of phosphate were made based on the methodology described for pH gradient plates ^(107)^. First 35 mL of molten MOPS agar at one end of the gradient was pipetted into a 10 cm square petri dish (Corning, BP124-05) that rested in a custom 3D-printed prop that held the plate slanted at a 30° angle. Once the bottom layer had solidified, the plate was laid flat and 35 mL of molten medium agar containing the second desired concentration on the gradient was poured atop. BCIP was added as described above.

### Calculating alkaline phosphatase enzymatic activity

A colorimetric assay using the substrate p-Nitrophenyl phosphate (PNPP) (New England Biolabs) was used to quantify alkaline phosphatase activity in colonies grown on MOPS agar plates with no added BCIP using the inoculation regime described above. After 12- 24 h incubation, the colonies were scraped from the agar with a pipette tip and resuspended in 1 mL 10 mM Tris-HCL buffer at pH 8. Fifty µL of the cell suspension was mixed with 50 µL of room temperature 1-step PNPP solution or 50 µL of Tris-HCL buffer at pH 8 and incubated for 1 h. A_405_ and A_600_ were recorded at t = 0 min and t = 60 min. AP activity was calculated using the equation 1000 x (ΔA_405_/(time (min) x A_600_)) where ΔA_405_ = (A_405_PNPP – A_405_NoPNPP)_t60_ – (A_405_PNPP – A_405_NoPNPP)_t0_.

### NanoString analysis

Total RNA was collected from *P. aeruginosa* colony biofilms grown on MOPS agar with 0.7 mM K_2_HPO_4_/KH_2_PO_4_. Bacteria were grown overnight in LB then sub-cultured in LB until OD_600_ = 0.5 before 5 µL were spotted on MOPS agar and incubated at 37 °C for 12 h. Cells were then scraped from the agar and resuspended in 200 µL Tris-EDTA buffer. RNA was extracted using the Qiagen RNeasy kit (Qiagen 74004) using the protocol provided by the manufacturer. Samples were not treated with DNAse. NanoString analysis of 80 ng of isolated RNA using codeset PaV5 ^(27)^ was performed as previously reported ^(108)^. Counts were normalized to the geometric mean of five housekeeping genes (*ppiD*, *rpoD*, *soj*, *dnaN*, *pepP*, *dapF*). Normalized counts were used for heatmap construction.

### Growth curves

*P. aeruginosa* was grown in LB medium for 16 h and then sub-cultured to a starting OD_600_ = 0.05 in 5 mL of MOPS minimal medium + 16.7 mM glucose in glass culture tubes. OD_600_ was measured using a Spectronic 20D+.

### Statistical analysis

Statistical analyses were performed using GraphPad Prism version 10.1.1 for macOS.

## Acknowledgments

The research reported in this publication was supported by grants from the Cystic Fibrosis Foundation, GREEN19G0 and STANTO19R0, and the National Institutes of Health (NIH), NHLBI T32HL134598 (A.C.) and NIDDK P30-DK117469 (Dartmouth Cystic Fibrosis Research Center or DartCF). Additional support came from bioMT (NIGMS P20GM113132) and the Dartmouth Molecular Biology Shared Resource (NCI 5P30CA023108).

## Notes

### Competing Interest Statement

The authors have declared no competing interest.

### Summary of Updates

We revised the text for clarity and removed Figure 6.

