## Supplementary material for "Loss of LasR function leads to decreased repression of *Pseudomonas aeruginosa* PhoB activity at physiological phosphate concentrations": Fig. S1, S2 and Table S1

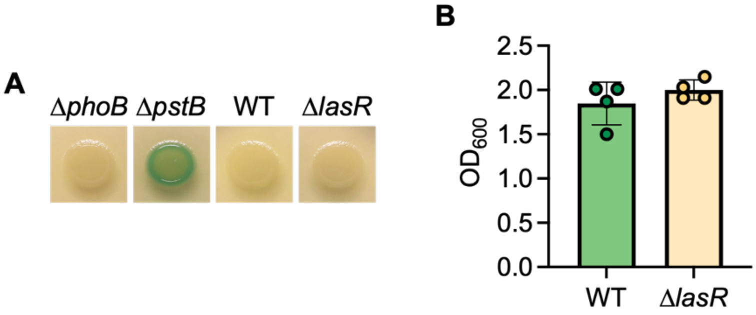


**Fig S1. *P. aeruginosa* wild type and Δ*lasR* strains show no PhoB activity on LB and grow similarly in MOPS with 0.7 mM Pi. A)** Colony biofilms of WT, Δ*lasR*, Δ*phoB* and Δ*pstB* were grown on LB agar with 60 µg/mL BCIP. Similar results were obtained in three replicate experiments and a representative image is shown above. **B)** *P. aeruginosa* was grown in MOPS liquid medium with 0.7 mM Pi. Data analyzed using an unpaired, two-tailed t-test ($p = 0.297$, $n = 4$).

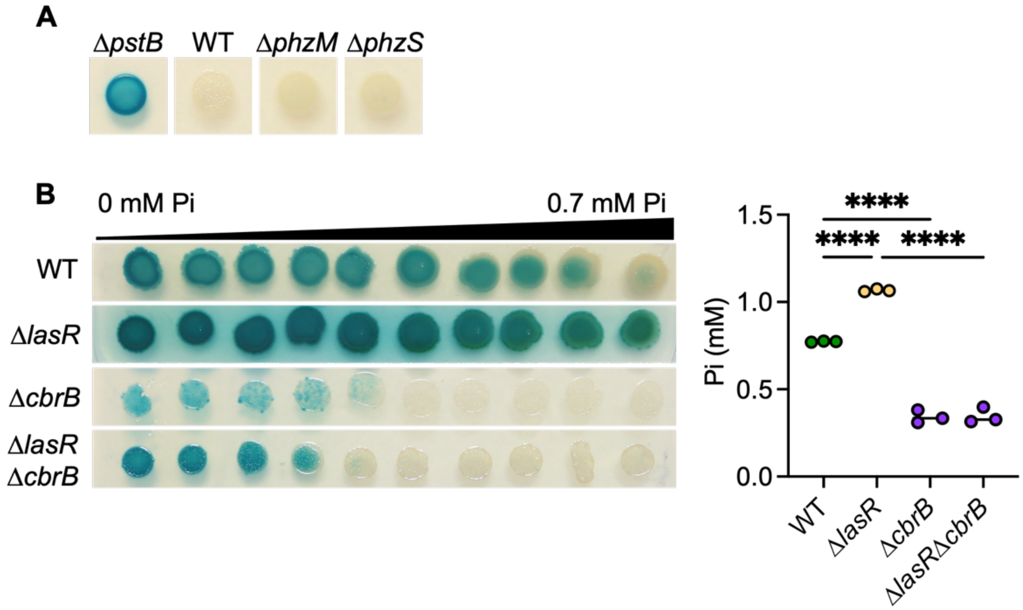


**Fig S2. Impacts of phenazine biosynthetic genes and CbrB on PhoB activity. A)** Colony biofilms of WT, Δ*pstB*, Δ*phzM* and Δ*phzS* were grown on MOPS agar with 0.7 mM Pi and 60 μg/mL BCIP. Similar results were obtained in three replicate experiments and a representative image is shown above. **B)** *P. aeruginosa* was grown on a plate with a gradient of Pi. The average concentration of Pi that inhibits AP activity is graphed to the right (n = 3). Data analyzed using a one-way ANOVA. Asterisks denote significance ($p \leq 0.0001$ = ****).

**Supplemental Table 1. Strains and plasmids used in this study.**

| Strain | Strain ID | Description | Source |
|---|---|---|---|
| *P. aeruginosa* | | | |
| PA14 WT | DH122 | Laboratory reference strain | (1) |
| PA14 Δ*lasR* | DH164 | DH122 with in-frame deletion of *lasR* | (2) |
| PA14 Δ*lasR* + *lasR* | DH3549 | DH164 with complementation of at the native locus | (3) |
| PA14 Δ*phoB* | DH3284 | DH122 with in-frame deletion of *phoB* | (4) |
| PA14 Δ*lasR*Δ*phoB* | DH4264 | DH3284 with in-frame deletion of *lasR* | This study |
| PA14 Δ*pstB* | DH3601 | DH122 with in-frame deletion of *pstB* | (4) |
| PA14 Δ*lasR*Δ*phoR* | DH4255 | In-frame deletions of *phoR* and *lasR* | This study |
| Clinical Isolate Pair 1 (LasR+) | DH1133 | Clinical strain AMT0047-2. CF lung isolate with functional LasR. | (5) |
| Clinical Isolate Pair 1 (LasR-) | DH1132 | Clinical strain AMT0047-3. CF lung isolate with loss-of-function mutation to *lasR*. | (5) |
| Clinical Isolate Pair 2 (LasR+) | DH2417 | Clinical strain NC-AMT0101-2. CF lung isolate with functional LasR. | (5) |
| Clinical Isolate Pair 2 (LasR-) | DH2415 | Clinical strain NC-AMT0101-1. CF lung isolate with loss-of-function mutation to *lasR.* | (5) |
| DH2590 | DH2590 | Clinical isolate from ocular infection with I215S mutation to *lasR.* | (6) |
| DH2590 + *lasR* | DH2743 | Clinical Isolate DH2590 with complementation of functional *lasR* at native locus | (3) |
| PA14 Δ*rhlR* | DH2742 | DH122 with in-frame deletion of *rhlR* | (7) |
| PA14 Δ*lasR*Δ*rhlR* | DH2944 | DH164 with in-frame deletion of *rhlR* | (7) |
| PA14 Δ*pqsR* | DH1110 | DH122 with in-frame deletion of *pqsR* | (8) |
| PA14 Δ*lasR*Δ*pqsR* | DH1111 | DH164 with in-frame deletion of *pqsR* | (9) |
| PA14 Δ*phz1* | DH1728 | DH122 with in-frame deletion of *phzA1-G1* operon | (10) |
| PA14 Δ*phz2* | DH1735 | DH122 with in-frame deletion of *phzA2-G2* operon | (10) |
| PA14 Δ*phz* | DH933 | PA14 with in-frame deletions of both *phzA1-G1* and *phzA2-G2* operons | (10) |
| PA14 Δ*lasR*Δ*phz* | SMC9422 | In-frame deletion of *phzA1-G1*, *phzA2-G2* and *lasR* genes | (11) |
| PA14 Δ*cbrB* | DH3920 | DH122 with in-frame deletion of *cbrB* | (12) |

| | | | |
|---|---|---|---|
| PA14 Δ*lasR*Δ*cbrB* | DH3924 | DH164 with in-frame deletion of *cbrB* | (12) |
| PA14 Δ*lasR*Δ*cbrB* + *cbrB* | DH3925 | DH3924 with complementation of *cbrB* at native locus | (12) |
| PA14 Δ*crc* | DH3737 | DH122 with in-frame deletion of *crc* | (12) |
| PA14 Δ*crc* + *crc* | DH3738 | DH3737 with complementation of *crc* at native locus | (12) |
| PA14 Δ*lasR*Δ*crc* | DH3927 | DH164 with in-frame deletion of *crc* | (12) |
| PA14 Δ*lasR*Δ*cbrB*Δ*crc* | DH3926 | DH3924 with in-frame deletion of *crc* | (12) |
| <u>*C. albicans*</u> | | | |
| CAF2 WT | DH48 | Laboratory reference strain | (13) |
| <u>*E. coli*</u> | | | |
| S17 λpir | DH71 | Used as a conjugation partner for introducing pMQ30 and GH121-based plasmids | |
| DH5α | DH51 | Used to store/replicate plasmids | Invitrogen |
| <u>Plasmids</u> | | | |
| pMQ30 EV | DH962 | Allelic replacement vector for use in yeast cloning, GmR | (14) |
| pMQ30_*lasR*_KO | DH2918 | *lasR* in frame deletion construction, GmR | (15) |
| pMQ30_*lasR* | DH3548 | *lasR* in frame complementation construct, GmR | (16) |
